# Evolution and Regulation of Decoupled Dimorphic Scaling Patterns in Ants

**DOI:** 10.1101/2025.08.25.672173

**Authors:** Erica Vong, Shannon Parisien, Helene Orfali, Rajendhran Rajakumar

## Abstract

Allometric evolution, changing how morphological traits scale disproportionately to body size, has fuelled adaptive radiations. In ants, allometry has facilitated the repeated evolution of a worker-soldier caste system. Here we reveal an unexpected scaling pattern, and its superorganismal regulation, within the hyperdiverse genus *Pheidole*. We discovered that antennal sizing between the small worker and the big soldier is identical, lacking inter-caste scaling (antennae-body scaling is decoupled) yet retaining intra-caste scaling. Furthermore, we have pinpointed the evolutionary origin of this decoupling. Finally, manipulations of social environment, developmental hormones, and interorgan signalling influences the modularity-integration of trait-body covariation, uncovering head-to-body decoupling and the production of novel nanosoldiers. Collectively, this challenges our previous understandings of trait covariation, modularity, and their regulation, from individual to society and across species.

## Main Text

Allometry, the scaling of morphological trait size to final body size, has fascinated the arts and sciences since the Renaissance (*1-12*). The evolution of allometry has generated an array of morphological novelties and has promoted adaptive radiations across the animal kingdom from the elongated legs of water striders and the beaks of Darwin’s finches, to the jaws of cichlids and the soldier caste in ants (*2, 13-15*). Variation in allometry can occur at the developmental-, individual-, and species-level (*15*). A proportional increase in trait size with body size is known as isometry, while a disproportionate increase or decrease in trait size with body size is known as hyperallometry and hypoallometry, respectively (*16*). Unexpectedly, we found a trait-body scaling relationship that does not conform to previously described allometric patterns, where the evolution of the soldier caste in *Pheidole* has facilitated a decoupled and scale-free exploration of otherwise forbidden morphospace.

The hyperdiverse genus *Pheidole* has a complex dimorphic worker-soldier caste system including small minor workers and big soldiers (*17*). This dimorphism is known to be regulated by a nutrition-dependent juvenile hormone (JH) developmental switch (*18-20)*, where high levels of JH induces bipotential larvae to determine their soldier identity – individuals with disproportionately larger heads, here illustrated with *Pheidole dentata* (Fig. 1A, B). Across the ants, patterns of allometry can be found for various morphological traits when compared to body size, especially the caste-defining large head of soldiers *(21)*. To investigate these allometric traits in ants, a few traits have traditionally been used as proxies for body size because they scale with body size, including the first antennal segment called the scape, as well as the width of the pronotum, and length of the hindleg tibia (*22-24*). Unexpectedly, in *Pheidole dentata*, we discovered a pattern that does not conform to previously described allometry: the mean and range of scape length variation shows a lack of inter-caste scaling when compared to head size, which is the caste-defining metric in *Pheidole* (Fig. 1C, E). When examining head size as compared to other known proxies for body size such as pronotum width and hindleg tibia length, inter-caste scaling is as expected (Fig. 1D & fig. S1A). In contrast, scape length compared to both pronotum width and hindleg tibia length comparisons demonstrate the lack of scaling to body size: scape length is indistinguishable and completely overlaps between workers and soldiers (Fig. 1E, fig. SlB, C). Furthermore, in the context of soldier body size, the scape is relatively far smaller than the relative sizing found in workers (Fig. 1F). Surprisingly, the lack of inter-caste scaling occurs despite there being scaling within each caste (Fig. 1C & fig. S1B,C). This suggests that the scape is coupled to body size within caste, but decoupled from body size between castes. Finally, when looking at the whole antennae as compared to head and proxies for body size, we observe the same presence of intra-caste scaling yet absence of inter-caste scaling as with the scape (fig. S1D). The scape is a morphological synapomorphy in ants that gives ant antennae their characteristic elbow shape, essential for individuals to perceive physical and chemical communications within and outside their colony (*25-27*). Collectively, the scape, a hallmark trait for social interactions in ants, is scale-free between workers and soldiers in *P. dentata*.

**Fig. 1.**
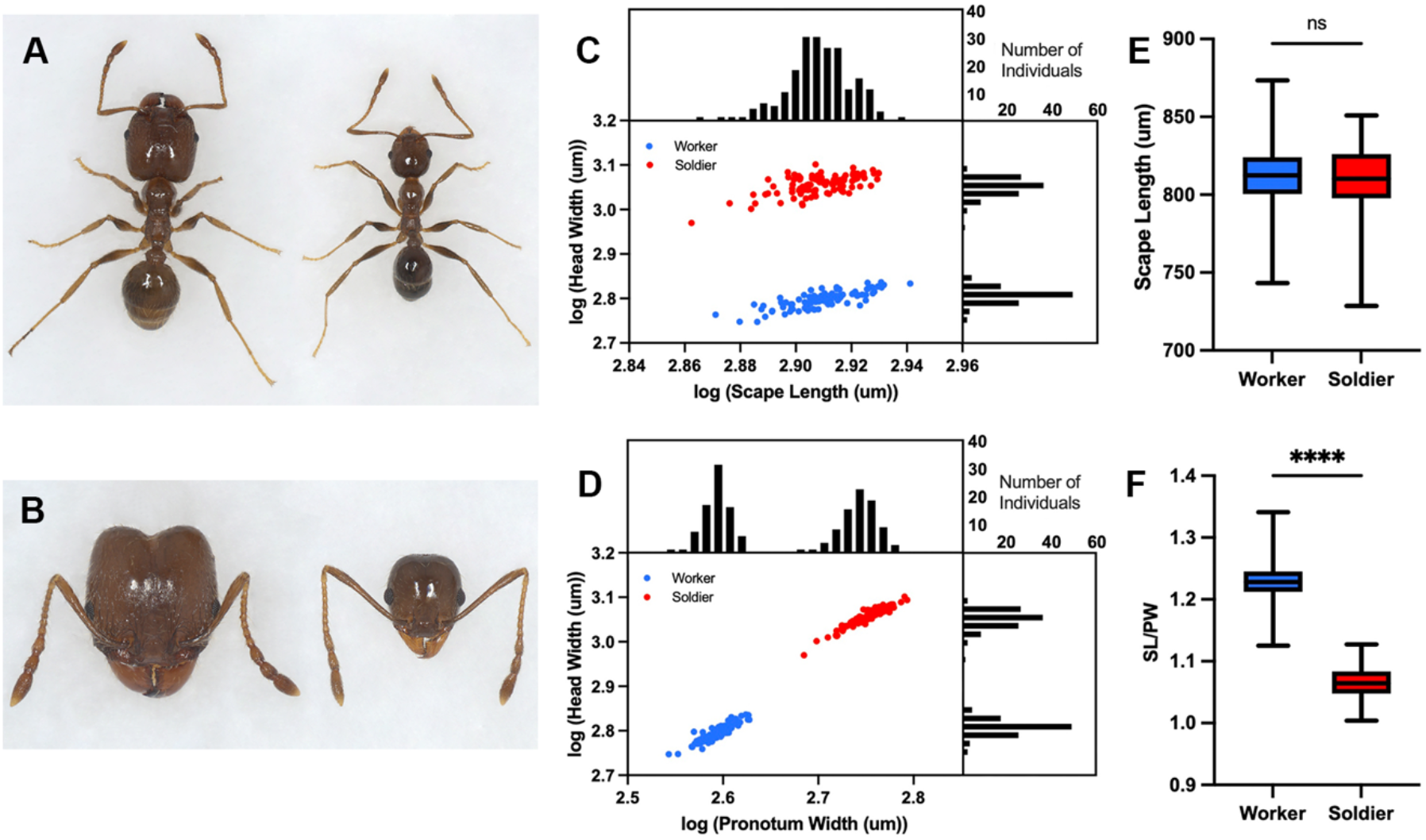
*Pheidole dentata* wild-type comparison of scaling and scale-free traits between workers and soldiers. **(A)** Larger soldier (left) and smaller worker (right) and the **(B)** larger heads in soldier compared to workers. Allometry log-log plots of **(C)** head width to scape length (n = 100) and **(D)** head width to pronotum width (n = 100), both graphs include frequency plots for number of individuals that corresponds to the respective X-axis (above each graph) and that corresponds to the respective Y-axis (on the right of each graph). **(E)** Box plot comparing scape length between workers (n = 100) and soldiers (n = 100) with median, interquartile range (bars), and minimum and maximum values (whiskers); unpaired two-tailed t-test, P > 0.05 and non-significant. **(F)** Box plot comparing scape length: pronotum width ratio between workers (n = 100) and soldiers (n = 100) with median, interquartile range (bars), and minimum and maximum values (whiskers); unpaired two-tailed t-test, ****P < 0.001.

To investigate whether the scale-free nature of the scape found in *P. dentata* occurs across the hyperdiverse genus *Pheidole*, we analyzed morphological measurements from 565 species of *Pheidole* from *Pheidole in the New World (17)*. First, we compared the scape to head width (caste identity) and scape to pronotum width (body size) across *Pheidole* and found that the total variation in scape sizes completely overlaps when looking across workers and soldiers of *Pheidole* (Fig. 2A, B). However, how worker and soldier scapes scale to one another within a species is unclear. We therefore developed a Dimorphic Scape Ratio (DSR), which is the ratio between the soldier scape to worker scape. Remarkably we find a normal distribution of DSRs, where the mode of the distribution is DSR=1, yet there were species that exhibited both DSR values > or < 1 (Fig. 2C). This is in stark contrast to the Dimorphic Head Ratio (DHR), where the mode of the distribution is DHR = 2, yet not a single species is less than 1.4 (fig. S2). Taken together, these data suggest that the decoupled nature of scape scaling between castes is the common state in *Pheidole* (Fig. 2C).

**Fig. 2.**
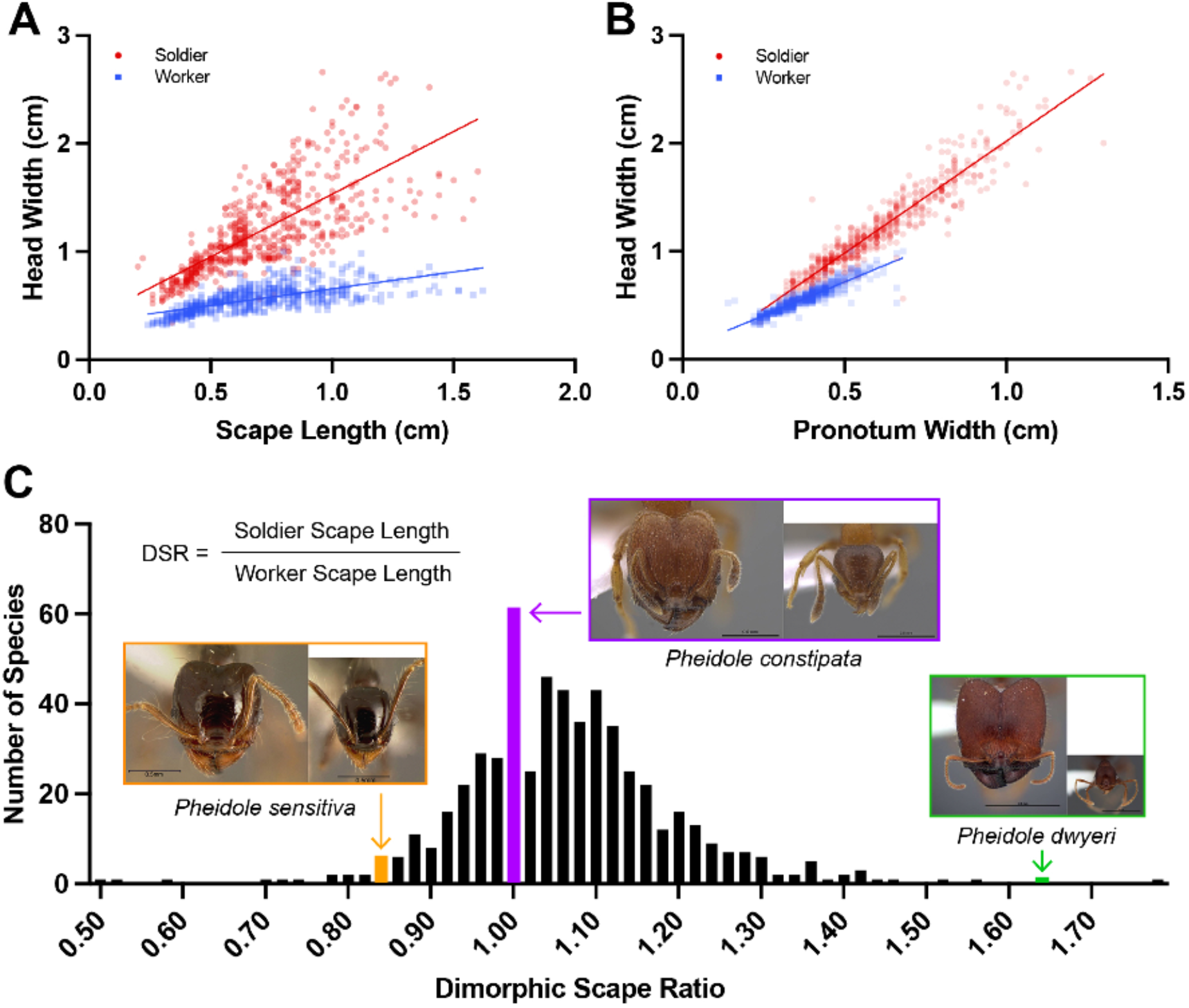
*Pheidole* genus analysis of genus wide allometry and Dimorphic Scape Ratio. **(A)** Head width versus scape length and **(B)** head width versus pronotum width for workers and soldiers using morphological measurements of 565 species collected from *Pheidole in the New World* (*17*). **(C)** Frequency of the Dimorphic Scape Ratio (DSR) which represents the soldier:worker scape length for 565 *Pheidole* species.

To determine whether the scape’s scale-free pattern exists in other ant species or if this is unique to *Pheidole*, we analyzed the scape-to-caste identity relationship by sampling across the ants, including 14 genera from 3 major subfamilies for species with a worker-soldier caste system and for related species with only workers (Fig. 3). In species that lacked a soldier caste we found that scape length scaled with body size. Similarly, in species that have independently evolved a worker-soldier caste system, we found that the majority of species have scapes that scale with body size, at both the intra-caste and inter-caste levels. Exceptionally, we discovered that *Cephalotes varians*, a species with a complex worker-soldier caste system, that is closely related to *Pheidole*, also have scape lengths that are decoupled from body size and caste identity. Taken together these data suggest that the origin of this scale-free pattern can be found at the common ancestor of *Pheidole* and *Cephalotes*.

**Fig. 3.**
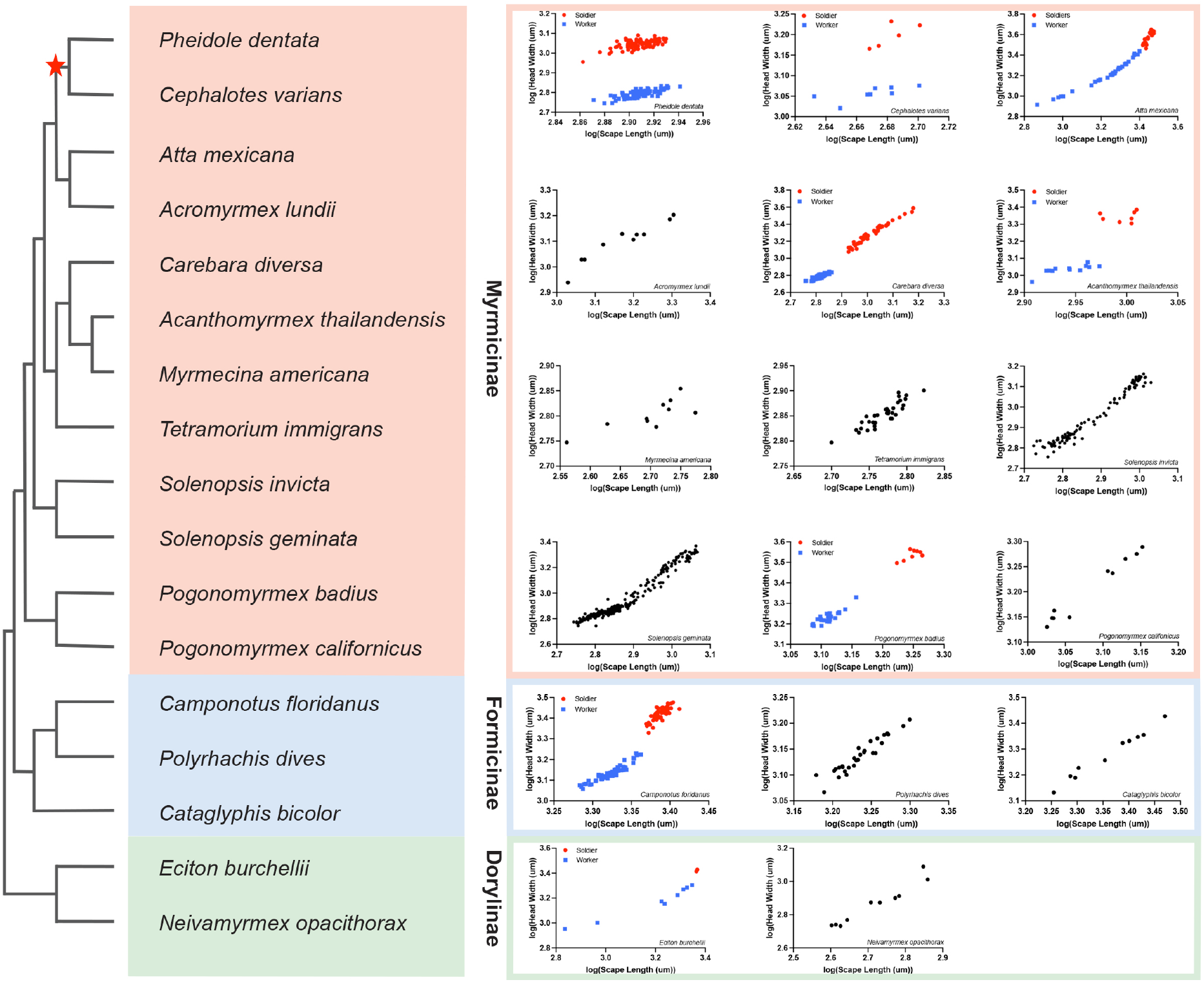
Phylogenetic Tree of Ant Species Grouped by Subfamilies with Head Width to Scape Length Allometry Pinpoints Origin of Decoupling. Evolutionary relationships and lineage divergence from 16 ant species categorized into their respective subfamilies: Myrmicinae (Beige; *Acanthomyrmex thailandensis, Acromyrmex lundii, Atta mexicana, Carabera diversa, Cephalotes varians, Myrmecina americana, Pogonomyrmex badius, Pogonomyrmex californicus, Polyrhachis dives, Solenopsis geminata, Solenopsis invicta, Tetramorium immigrans*), Formicinae (Blue; *Camponotus floridanus, Cataglyphis bicolor, Polyrhachis dives*), and Dorylinae (Green; *Eciton burchellii, Neivamyrmex opacithorax*). Head width to scape length allometry plots included where red represents soldiers, blue represents workers, and black represents workers in species lacking a morphological worker-soldier caste system. Star denotes the potential origin of decoupled traits.

The determination of caste identity is regulated at multiple levels of biological organization in *Pheidole* including the social/pheromonal and hormonal levels (20,*28-31*). It is known that nutrient-dependent juvenile hormone positively regulates soldier caste determination in *Pheidole* during a critical window of development *(18, 20, 31, 32*). Moreover, it is known that social demography regulates the colony ratio of soldiers to workers through an inhibitory pheromone produced by soldiers that negatively regulates soldier caste determination (*28, 30, 32*). Based on this, we next tested how these regulatory levels influence the decoupling of scape allometry from caste identity in *Pheidole*. We treated bipotential larvae with methoprene (JH analog) or acetone (control) and raised by either 100% workers or 100% soldiers. As compared to workers generated in the acetone control, JH-treated 100% worker-raised bipotent larvae developed into soldiers exhibiting head and scape variation as well as head-to-scape allometry as found naturally (Fig. 1C, Fig. 4A-C, fig. S3A). In the context of manipulating JH and the inhibitory pheromone (soldier-raised), we found that bipotent larvae exposed only to the inhibitory pheromone generated significantly smaller workers with disproportionately (different y-intercept) smaller scapes as compared to bipotent larvae exposed to both the inhibitory pheromone and treated with JH (Fig. 4D-F). While the workers generated from the combination of pheromone and JH are reflective of typical workers found in the colony (fig. S3B), the individuals generated from exposure to the inhibitory pheromone without JH were smaller in scape, head, and body size as that found naturally (Fig. 4E, F, fig. S3B, C). Therefore, in the context of both soldier identity being induced by JH (Fig. 4A) and soldier identity being suppressed by soldier-skewed caste ratio (Fig. 4D), scape pattern remains decoupled from caste-identity.

**Figure 4.**
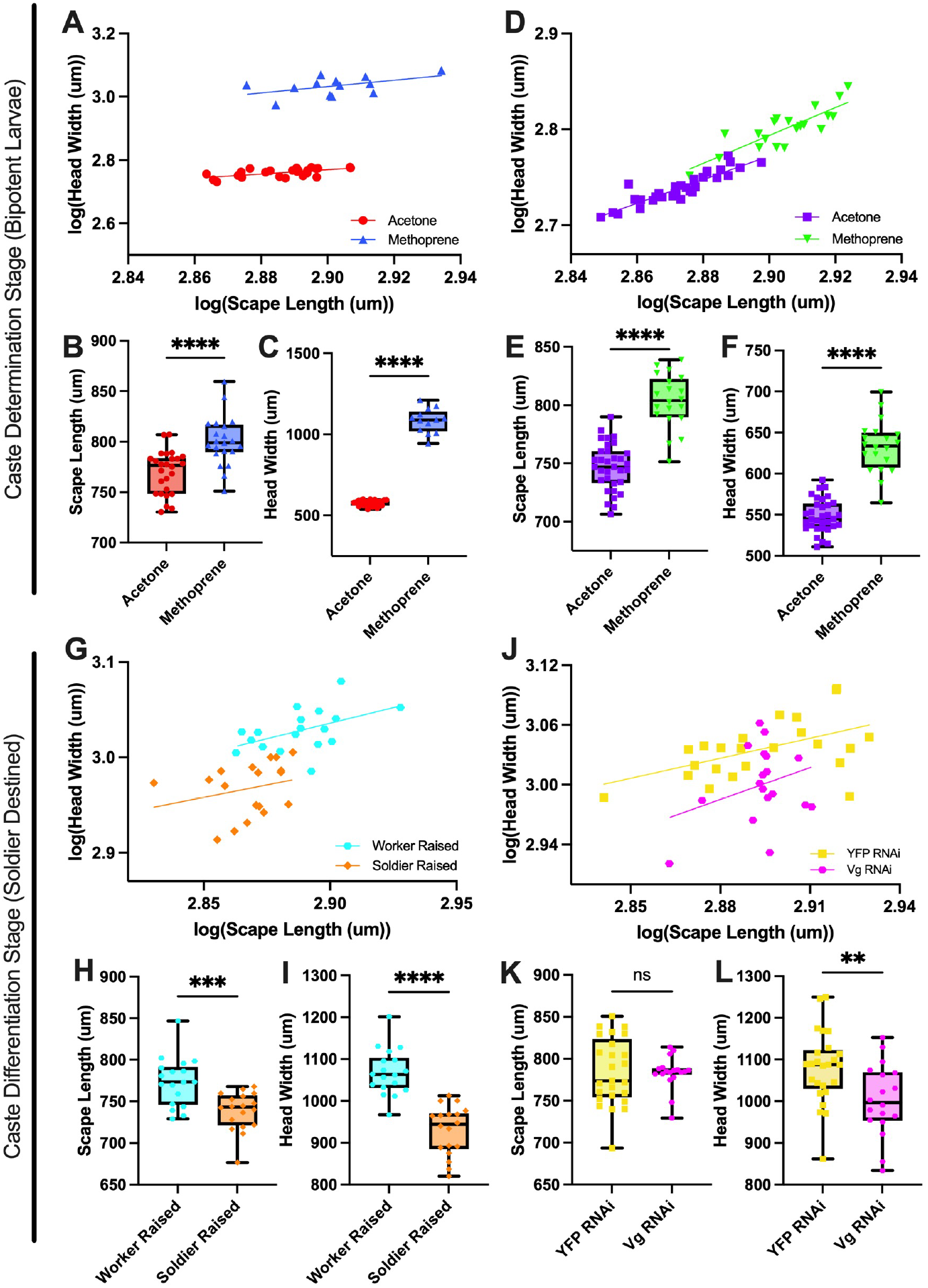
Social and Hormonal Manipulations on *Pheidole dentata* at the Caste Determination and Caste Differentiation Developmental Timepoints. Bipotential larvae (caste determination) that are worker raised: **(A)** log-log plot of head width versus scape length for acetone control (n = 26) and methoprene-treated (n = 13) individuals (ANCOVA: slope P>0.05; y-intercept, F=1409, df=36, ****P<0.0001) **(B)** significant differences in their scape length (t=4.661, df=45, ****P < 0.0001) and **(C)** in their head width (t=33.24, df=37, ****P < 0.0001). Bipotential larvae that were soldier raised: **(D)** log-log plot of head width versus scape length for acetone control (n = 33) and methoprene-treated (n = 20) individuals (ANCOVA slope P>0.05; y-intercept, F=21.37, df=50, ****P<0.0001) **(E)** significant differences in their scape length (t=9.353, df=51, ****P < 0.0001) and **(F)** in their head width (t=11.79, df=51, ****P < 0.0001). For experiments on soldier-destined (caste differentiation) **(G)** log-log plot of head width versus scape length for worker-raised (n = 18) and soldier-raised (n = 18) individuals (ANCOVA slope P>0.05; y-intercept, F=25.58, df=33, ****P<0.0001) **(H)** significant scape length (t=4.023, df=34, ***P = 0.0003) and **(I)** head width (t=7.341, df=34, ****P < 0.0001). Soldier-destined **J)** log-log of head width to scape length for *vestigal* RNAi (n = 18) and YFP control (n = 25) (ANCOVA slope P>0.05; y-intercept, F=13.86, df=39, ***P<0.001) **K)** non-significantly different scape length (P=0.8983) and **L)** significantly different head widths (t=2.898, df=41, **P<0.006).

Caste differentiation, is a critical developmental period following caste determination where caste-specific scaling of morphological characters unfolds. Previously it has been shown that the scaling of developing individuals, for which caste identity has been acquired (soldier-destined), can be impacted by social interactions and interorgan signaling *(32)*. Specifically, the scaling of soldier-destined individuals is altered when they are raised in a soldier inhibitory environment and when interorgan communication is perturbed (*32*). To test the effects of social interactions on scape scaling, we first analyzed soldier-destined individuals raised by soldiers as compared to raised by workers. The inhibitory pheromone generated significantly smaller individuals with disproportionately (different y-intercept) smaller scapes as compared to those raised by workers (Fig. 4G-I, fig. S3D, E). Surprisingly, these individuals have soldier-sized heads yet have the same sized bodies as wildtype workers (fig. S3D, E). Therefore, caste-identity (giant versus small head size) is no longer coupled to body size, generating individuals, which we here describe as ‘nanosoldiers’ that have novel extreme head- and scape-to-body allometry (fig. S3D-F). To test the effects of interorgan signaling on scape scaling, we next perturbed the rudimentary wing disc of soldier-destined larvae, a tissue known to regulate head-to-body allometry and soldier identity by acting as a signaling center through interorgan communication *(32)*. This was done by performing RNAi (RNA interference) of *vestigial*, previously shown to halt soldier rudimentary wing disc growth and soldier-specific development in *Pheidole (32)*. Perturbing the rudimentary wing disc of soldier-destined larvae generated significantly smaller individuals, with significantly smaller heads, including nanosoldiers (Fig. 4J-L, fig. S3G, H). However, unlike with the soldier inhibition experiments, the average scape size of these individuals were not significantly different compared to controls (Fig. 4K). Surprisingly, while soldier-destined controls exhibited scape variation around the mean, soldier-destined individuals treated with *vestigial* RNAi completely lacked scape variation (i.e.: variable head and body size, yet invariable scape; Fig. 4J-L, fig. S3G, H). Taken together, social and interorgan communication can influence both scape and head allometric coupling and decoupling during caste differentiation. Collectively, manipulating pheromones, growth hormones, and allometric signaling tissues can both generate a quantitative shift in pre-existing intra- and inter-caste allometry, and generate novel inter-caste variation.

Despite the ubiquity of coupled scape-to-body allometry across the ants, we have found that decoupled worker-soldier scape-to-body allometry is a common feature to the hyperdiverse genus *Pheidole*. Furthermore, we found that in parallel to *Pheidole, Cephalotes varians* have decoupled worker-soldier scape-to-body allometry. Notably, *Pheidole* is sister to *Cephalotes* + *Procryptocerus* (PPC clade; *33, 34)*; *Procryptocerus* are ants that lack a soldier caste (*35*). Interestingly, *Procryptocerus* workers and queens differ in body size (*35, 36*), yet they are ambiguous with regards to their head size *(35)*. If this decoupling is found to be prevalent in *Cephalotes*, it is possible that traits decoupled from body size between castes is the ancestral state of these three genera and that these clades have the developmental potential to decouple trait variation and scaling from size and caste identity.

Many insects rely on their antennae as an organ for tactile exploration to navigate their physical environment and for detecting odors of a variety of things including food and conspecifics (*27, 37*). In ants, tactile information is perceived by mechanoreceptors and odors by chemoreceptors located on the antennal sensilla (*38, 39*). Ants are social insects that use their antennae to physically and chemically communicate information on a societal level to cooperatively hunt prey, defend against predators, perform colony tasks, and even to control the caste demographic of the colony (*40, 41*). A key synapomorphy of ants includes the elongated first antennal segment, the scape, which connects via an elbow-like joint to the flagellum which houses the majority of the sensilla (*25, 42*). The lengthened scape allows for the ants to increase the physical range they can sense; it has also been shown to actively change its angle to control the position of the sensilla-dense tip of the antenna when exploring new environments *(27)*.

Across the ants, our data shows that antennae scale to an individual’s body size. This is in contrast to what we have found in *Pheidole* where the adaptive role of the observed worker-soldier antennal decoupling remains unclear. We hypothesize that at the origin of the PPC clade, a novel developmental mechanism has evolved that provides the capacity for some traits to explore their morphospace in a caste-specific manner while other traits are constrained such that they are decoupled from caste-specific development. We additionally hypothesize that the shared worker-soldier variation range in scape length is a result of functional and structural constraints, and that the variation itself within each caste is a by-product of the environmental and hormonal influence on an individual’s genome, thereby their degree of developmental plasticity.

Superorganismal interactions in *Pheidole* include nutrition-dependent hormonal regulation and inhibitory pheromone regulation *(32)*. Together these interactions influence interorgan signaling enabling trait covariation and caste-specific evolution. Our study demonstrates that some traits have evolved developmental mechanisms decoupling them from the integrated covariation of caste-specific traits enabling the modular development of intra-caste trait variation independent of inter-caste variation. The striking degree of this modularity is moreso unexpected because the head and antennae share a common developmental origin (*43*), yet the head is hypersensitive to both soldier-specific circulating hormones and interorgan signals, whereas the antennae demonstrates robustness to these cues, buffering developing antennal cells from responding to soldier-specific signals and canalizing the phenotype at the inter-caste variation level (Fig. 5B). During caste determination, manipulating superorganismal conditions had no effect on the decoupled scape pattern we have found. Surprisingly, during soldier differentiation we found that exposing individuals to a soldier inhibition environment generated a nanosoldier caste that exhibited exaggeratedly large heads relative to their worker-sized bodies (fig. S3D); completely independent of body size variation. This indicates that following the soldier determination switchpoint, JH-dependent soldier-specific head growth is irreversible and proceeds, yet the soldier pheromone inhibits systemic growth and body sizing, generating nanosoldiers. Therefore, while developmentally there is a potential for head size to be decoupled from body size, we nevertheless do not find nanosoldiers in the wild indicating that there is selection against this morphospace rendering this unexplored variation forbidden. Furthermore, developing bipotential individuals exposed to the soldier inhibition environment generates nanoworkers with proportionally smaller scapes, which do not exist in nature (Fig. 4D, fig. S3B). We suggest these nanoworkers were generated due to the soldier inhibitory pheromone influencing growth pathways involved in intra-caste proportional sizing independent of both JH and rudimentary wing disc influences as workers lack any rudimentary wing disc (Fig. 5B) (*32*). When considering nanosoldiers in the context of large workers and nanoworkers, two unanticipated relationships between the head, body, and scape, emerge (Fig. 5A). First, nanosoldiers have larger head sizes than large workers (fig. S3D), while they have the same body size (fig. S3D) yet extremely smaller scapes (fig. S3F). This is surprising since *Pheidole* scape decoupling would predict that individuals that exhibit large worker body size to have a large scape, however nanosoldiers have recoupled their scape-to-body relationship across worker and soldier size ranges (fig. S3F). Second, nanosoldiers have larger body size and head size than nanoworkers, yet their scapes are indistinguishable (Fig. 5A, fig. S3I). In contrast, *P. dentata* queens have larger bodies with proportionally larger scapes suggesting that this species retains the developmental pathways (caste-specific developmental plasticity) necessary for integrated covariation coupling scape and body size despite these pathways being suppressed (non-plastic) in the worker-soldier developmental trajectory (fig. S4A-C). Altogether, the influence of superorganismal regulation, through inhibitory pheromones, on scaling and decoupling of traits depends on whether these molecules are acting during caste determination or caste differentiation.

**Figure 5.**
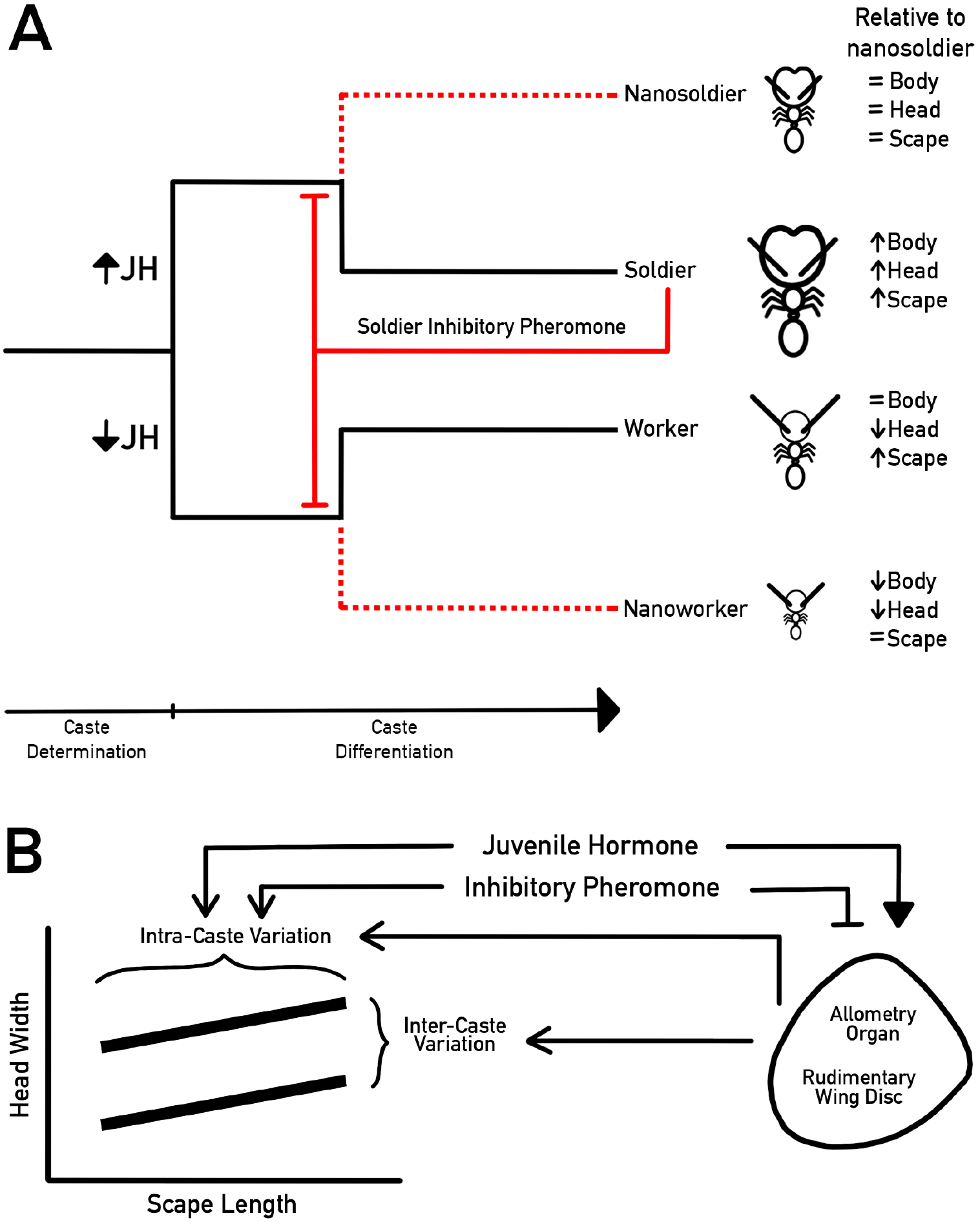
The interplay between developmental hormones, social pheromones, and inter-organ signaling on inter-caste determination and intra-caste differentiation. **A)** *Pheidole* worker-soldier developmental trajectories where JH mediates caste determination and the soldier inhibitory pheromone influences intra-caste variation during the early caste differentiation period generating nanoworker and nanosoldier variation. **B)** The allometry organ, hormones, and pheromones as sources of variation (arrow). The influence of juvenile hormone (arrowhead) and inhibitory pheromone (bar) on the allometry organ further contributes to intra- and inter-caste variation.

Perturbing rudimentary wing imaginal discs in soldier-destined larvae with *vestigial* RNAi led to the generation of individuals with both smaller bodies and heads, yet invariable scape lengths reflective of the average sized worker and soldier. This indicates that for the antenna, caste differentiation processes were arrested, generating a canalized phenotype where there is one sized scape for several head-to-body-sizes (Fig. 4J, L). We suggest this is possibly due to the loss of developmental plasticity that normally generates intra-caste variation independent of the developmental plasticity mechanisms that generates inter-caste variation. Therefore, the interorgan signaling soldier rudimentary wing disc, known to regulate caste identity and allometry, additionally integrates environmental cues to regulate intra-caste variation (Fig. 5B).

Our data shows that traits can be decoupled to varying degrees both naturally and artificially. Across allometry research, it is common to assume that larger individuals have larger traits whether it follows a hyper- or hypoallometric pattern. In *P. dentata* when analyzing how the scape varies with increases in body size, the data is nonsensical as there is no correlation with body size and therefore the antennae is scale-free. In contrast, only in the context of the biological castes, do we resolve the presence of trends that exists within a caste (intra-caste scaling) rather than between castes (inter-caste scaling). This therefore turns otherwise statistical noise into signal or maladaptive/neutral morphological variation into adaptive phenotypes. The absence of a trend occurring when knowledge of separate groups is lacking (grouped together), but the existence of trends when separate groups are considered, is known as Simpson’s paradox (*44, 45). P. dentata* provides a biological example for which this paradox can be resolved when considering the worker-soldier caste system. The structure of antennal-to-body allometry and variation observed would be maladaptive to an organism for which traits would normally need to scale to the body due to selection, yet by evolving two castes, this system composed of two caste-specific groups, have resolved this paradox and are independently adaptive for intra-caste antennal-to-body allometry. Likely the biological resolution of Simpson’s paradox for various traits has occurred throughout plants and animals whether by genetic differentiation (ex: sexual dimorphism) or developmental plasticity. Our work looking at the DSR and DHR of *Pheidole dentata*, raises the question of how dimorphic ratios across the body plan and across species have evolved. The evolution of the *Pheidole* worker-soldier caste system and both the modular nature and superorganismal regulation of development of the *Pheidole* caste system has facilitated decoupling and elaboration of morphology to a radical degree. More generally, increasing the number of modular plastic biological units in a system, whether gene, cell, organ, or caste can facilitate the exploration of novel trait variation and adaptive evolution.

## Acknowledgments

We thank L. Davis for help with ant collection; C. Jelley, M. Barkdull, and C. Moreau for help with access and training at the Cornell University Insect Collection; A. Rajakumar and Rajakumar Lab members for comments.

## Author contributions

Conceptualization and experimental design: E.V., R.R.

Experimentation: E.V., S.P., H.O., R.R.

Visualization and analysis: E.V., S.P., H.O., R.R.

Funding acquisition: E.V., R.R.

Supervision: R.R.

Writing – original draft: E.V., R.R.

Writing – review & editing: E.V., R.R.

## Data and materials availability

All data are available in the main text or the supplementary materials.

## Supplementary Materials

Materials and Methods

Figs. S1 to S5

References (1–48)

## Materials and Methods

### Animal Collection and Culturing

We collected queenright colonies of *Pheidole dentata* and *Camponotus floridanus* from Gainesville, Florida, USA. Ant colonies were maintained in lab in plastic boxes lined with fluon, which created a friction-less surface when dried to retain ants in their boxes. Boxes contained glass tubes filled with water and sugar water constrained with cotton. Colonies were fed a combination of mealworms, crickets, and Bhatkar-Whitcomb diet (*46*). Ant colonies were maintained at 25°C, 60% humidity, and 12h light:dark cycle.

### Adult Samples for Wild-Type Morphometrics

*Pheidole dentata, Camponotus floridanus, Carebara diversa, Acanthomyrmex thailandensis, Tetramorium immigrans*, and *Polyrhachis dives* are species where live adults were collected in lab, anesthetized, and stored in ethanol to be dissected for morphometrics. *Acromyrmex lundii, Myrmecina americana, Pogonomyrmex californicus, Cataglyphis bicolor, Eciton burchellii*, and *Neivamyrmex opacithorax* are species where data was collected from the Cornell University Insect Collection (CUIC) and a sample size of 10 for each of these species was obtained. To supplement museum samples, we further measured images from Antweb.org for species from the CUIC. *Atta mexicana* and *Cephalotes varians* are species whose adult samples were donated by Dominic Ouellette and Megan Barkdull, respectively. *Pogonomyrmex badius* adults (minor workers and soldiers) were collected in Gainesville, Florida, USA. *Solenopsis invicta and Solenopsis geminata* morphometric data is reanalyzed from (*47*). Scape data on 565 *Pheidole* species was extracted from that written and recorded in *Pheidole in the New World* (*17*) in addition to head width and pronotum width.

### Morphometrics and Statistical Analyses

We used a Zeiss Axio Zoom V16 stereomicroscope and ZEN Pro 3.2 Software to measure the larval length for the hormonal and soldier-destined experiments, in addition to all adult samples. Adults were measured in µm for head width, scape length, pronotum width, and hindleg tibia. At the CUIC, specimens were imaged using a Canon EOS D6 with a Canon MP-E 65mm Macro Photo lens. Images from the CUIC and Antweb.org were measured using ImageJ in mm for head width, scape length, and pronotum width. Values were log transformed and a simple linear regression was performed for each group and compared using an ANCOVA. We used an unpaired *t*-test to determine the difference between means of scape length, head width, and hindleg tibia. For the analysis across *Pheidole* using (*17*), 59 species were excluded for the genus-wide analysis of scape length due to missing values for either or both worker-soldier castes.

### Hormonal & Social Manipulations

We ectopically applied methoprene (CAT: 33375 ; Millipore Sigma), a synthetic analogue of JH, to the ventral side of *Pheidole dentata* larvae. To explore the effect of hormonal and social environment as well as their interplay, larvae were exposed to either inhibitory pheromone or hormone and their respective controls. For hormone it is methoprene-treated or control acetone-treated larvae, and for inhibitory pheromone, the social environment is composed of larvae that are raised by either 100% workers or 100% soldiers. We applied 1µL of methoprene dissolved in acetone at a concentration of 5mg/mL to larvae ranging from 1000-1500um in length, a size range prior to their determination into a minor worker or soldier. We apply 1uL of acetone as a control. We raised these methoprene or acetone-treated larvae in replicate boxes that contained a 2:1 adult-to-larva ratio.

To test the contribution of the social environment on developing soldiers, we collect larvae passed the worker-soldier switchpoint (soldier differentiation period). Specifically, we collect larvae ranging from 2000-2400um in length, a size range exceeding the terminal minor worker size but prior to terminal soldier size. These early soldier-destined developing individuals were then raised in a social environment of 100% soldiers. As a control, we raise larvae of this size range in a 100% worker environment.

### Cloning of *P.dentata vestigial* (*vg*)

Forward: 5’-TATCCTTACCTKTAYCARACCC-3’ and reverse: 5’-GTGTTCCTCSACYTCYGACG-3’ degenerate primers were used based on that previously done (*32*). *P. dentata* mRNA was isolated using TRIzol (CAT: 15596026; Invitrogen) from embryonic, larval, and pupal developmental stages. Developmental mRNA was then reverse transcribed to synthesize a cDNA library for subsequent amplification of *vg*. PCR amplificons were then ligated into a pGEM-T easy vector (CAT: A1360 ; Promega) and Sanger Sequenced at Genome Quebec. This 585bp *vg* fragment was subsequently used to synthesize double-stranded RNA (dsRNA) for RNAi.

### *vestigial* RNA interference (RNAi)

To RNAi knockdown *vestigial* in *Pheidole* with dsRNA in the same manner as it was done previously to disrupt the development of the soldier rudimentary wing disc (*32*), T7 flanked (uppercase) *vg* forward: 5’-TAATACGACTCACTATAGGG tatccttacctgtatcagaccc-3’ and reverse 5’-TAATACGACTCACTATAGGG gtgttcctccacctccgacg-3’ primers were used to design a 585bp fragment of dsRNA for RNAi (fig. S5). Primers were designed by adding the T7 promoter sequence upstream of species*-*specific primer sequence determined from the *P.dentata* cloned fragment. Purified *vg* amplicon was then used as a template transcribed using T7 polymerase (CAT: EP0113; ThermoFisher) then purified using MEGAclear Transcription Clean-up Kit (CAT: AM1908; Invitrogen) and finally, concentrated by ethanol precipitation. Soldier-determined larvae (2000-2400um larval length) were injected with either 5000ng/uL *vg* dsRNA or 5000ng/uL YFP (yellow fluorescent protein; as previously (*48*)) and raised by 100% workers with a 2:1 adult to larval ratio until the adult stage. Individuals were stored in 75% ethanol for dissection and morphometrics.

**Fig. S1.**
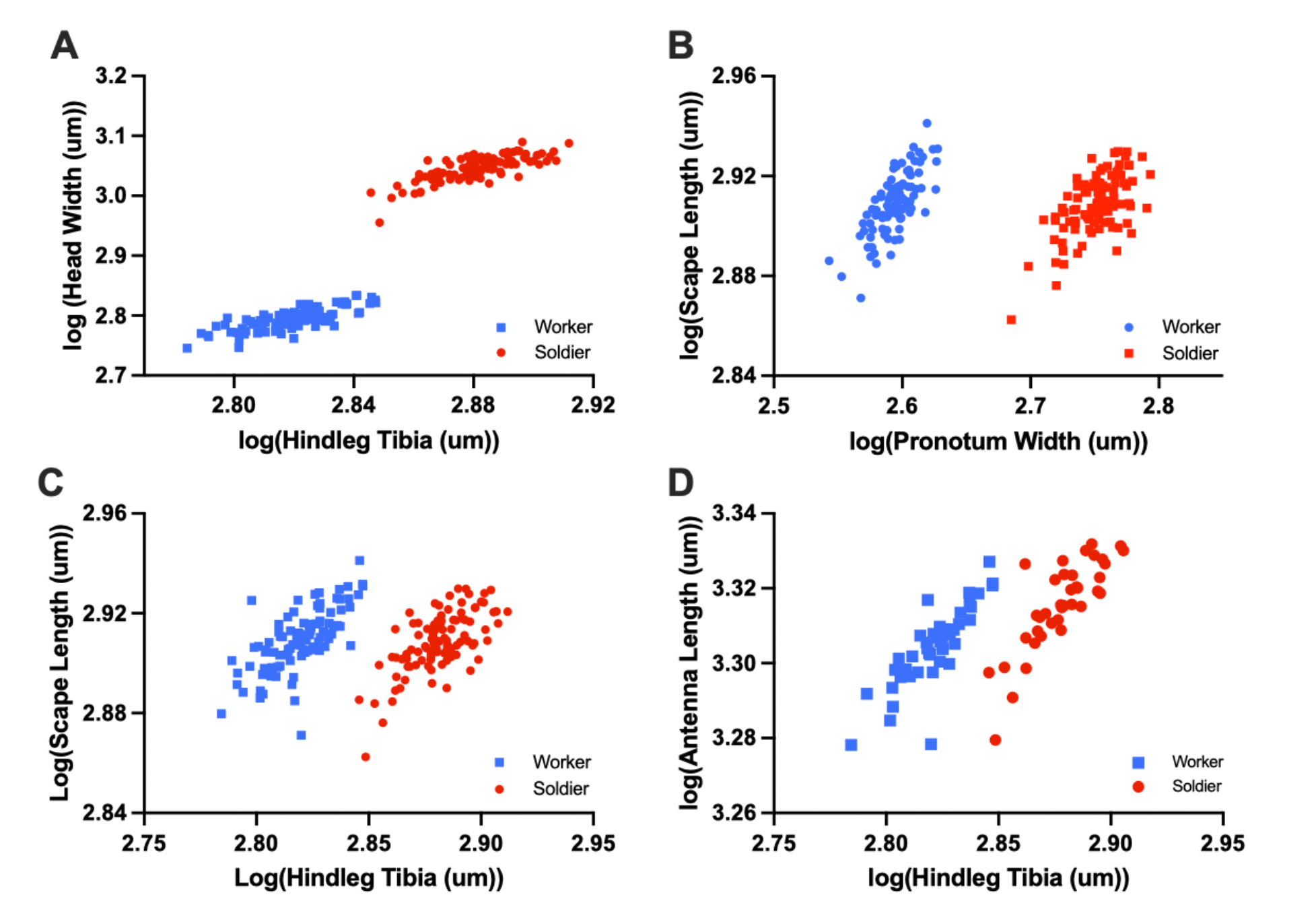
*P. dentata* allometry plots of log-log **A)** head width to hindleg tibia, **B)** scape length to pronotum width, **C)** scape length to hindleg tibia, and **D)** whole antenna length to hindleg tibia where workers are denoted in blue squares and soldiers in red circles.

**Fig. S2.**
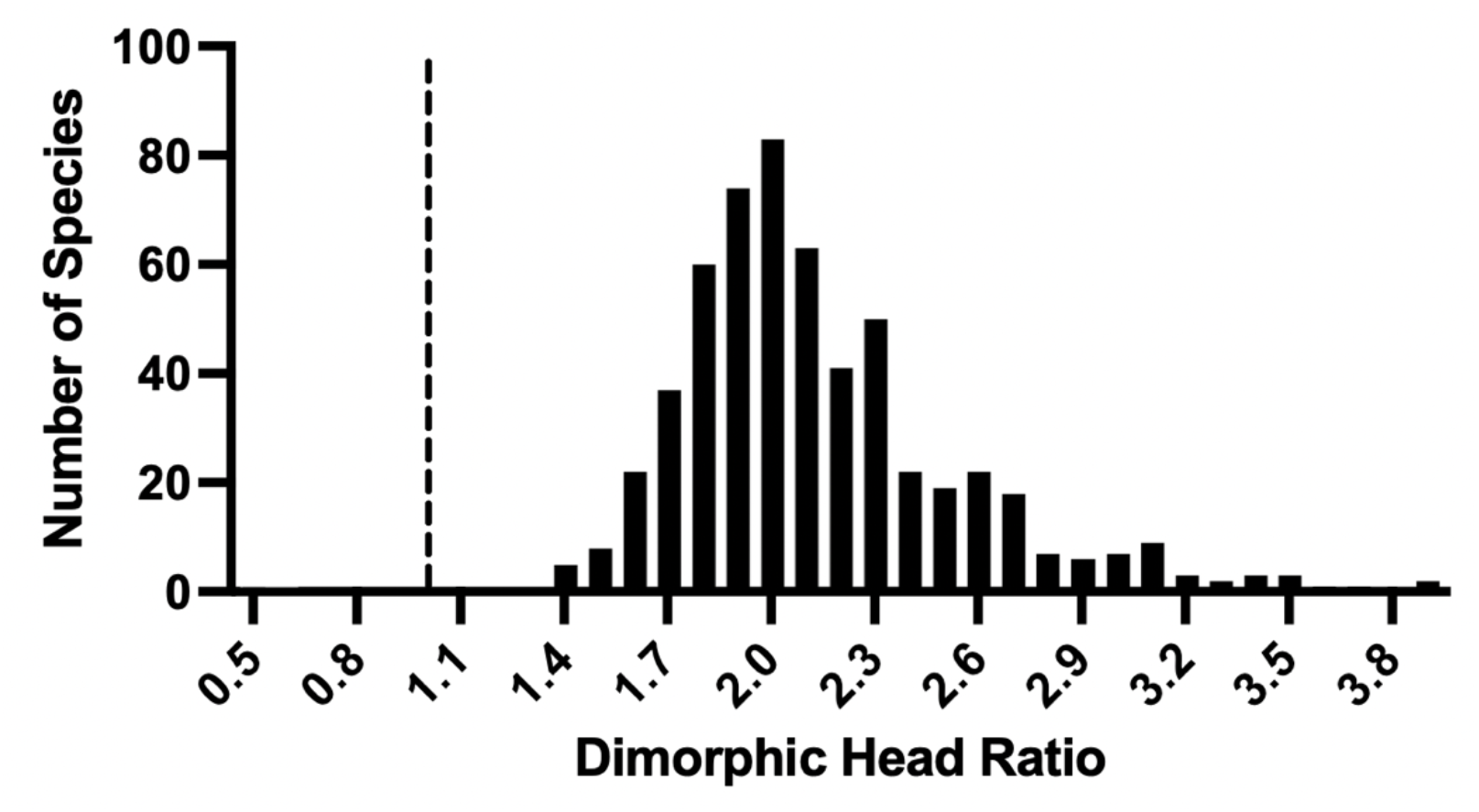
*Pheidole* genus analysis of the Dimorphic Head Ratio (DHR) which represents soldier:worker head width ratio for 565 *Pheidole* species from *Pheidole in the New World* (*17*). Dotted line denotes a DHR value of 1.

**Fig. S3.**
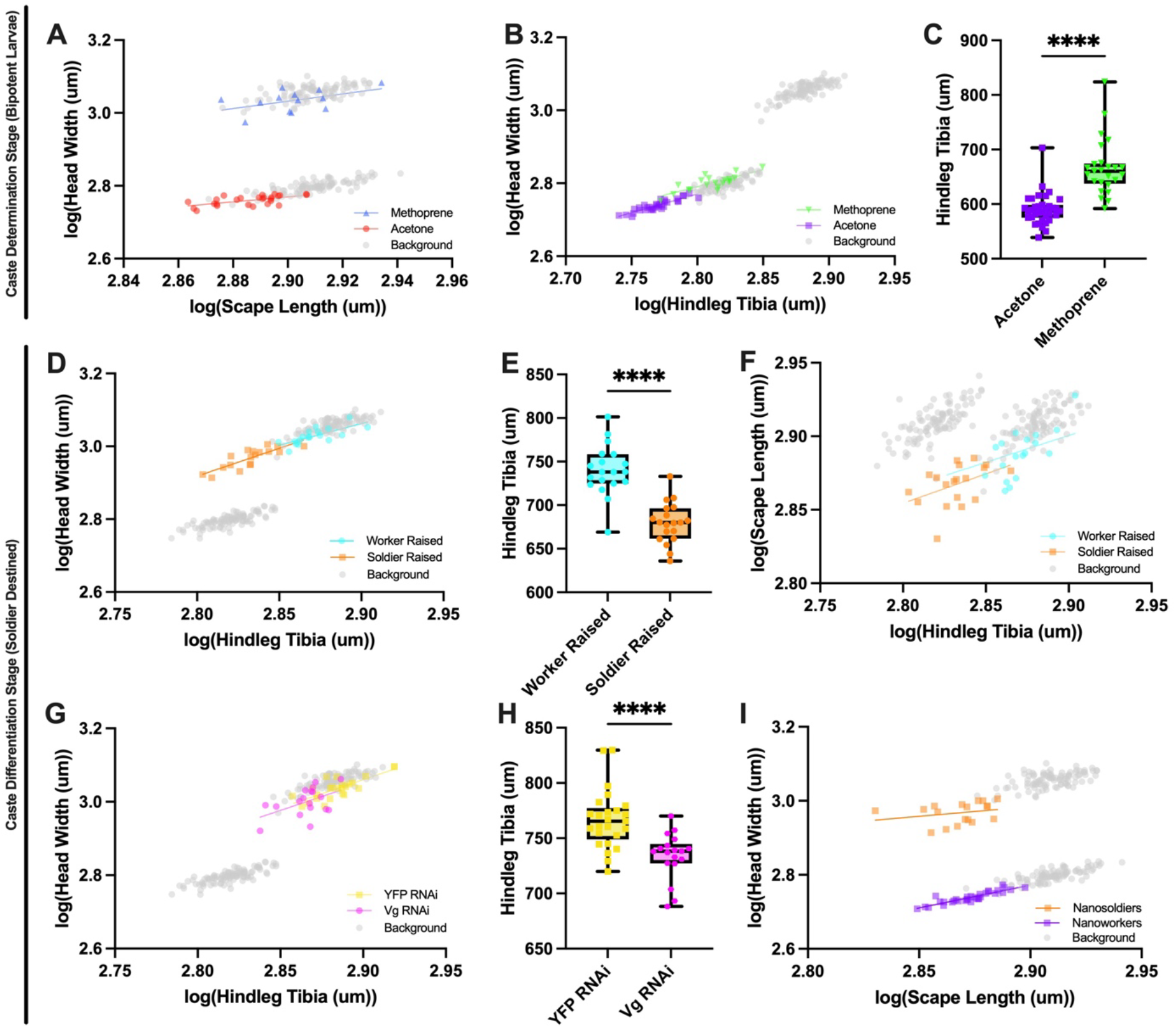
Social and hormonal manipulation for the caste determination stage using bipotent larvae with wildtype background variation. **A)** Bipotential larvae are raised by 100% workers treated with methoprene or acetone log-log plot of head width to hindleg tibia (ANCOVA slope P>0.05; y-intercept, F=1409, df=36, P<0.0001) with background variation. **B)** Bipotential larvae are raised by 100% soldiers treated with methoprene or acetone log-log plot of head width to hindleg tibia (ANCOVA slope P>0.05; y-intercept, F=21.40, df=50, P<0.0001) with background wildtype variation and **C)** significantly different hindleg tibias (t=7.364, df=58, P<0.0001). Social and tissue manipulation for the caste differentiation stage using soldier-destined larvae where **D)** 100% worker or 100% soldiers raised soldier-destined larvae log-log plot of head width to hindleg tibia (ANCOVA slope and y-intercept P>0.05) with background wildtype variation, **E)** difference in hindleg tibia length (t=7.119, df=36, P<0.0001), and **F)** log-log plot of scape length versus hindleg tibia (ANCOVA slope and y-intercept P>0.05) with background wildtype variation **G)** *Vestigal* knockdown in soldier-destined larvae log-log plot of head width to hindleg tibia (ANCOVA slope and y-intercept P>0.05) with background wildtype variation and **H)** significantly different hindleg tibias (t=4.351, df=41, P<0.0001). **I)** Log-log plot of head width compared to scape length between nanoworkers and nanosoldiers (ANCOVA slope P=0.0831; y-intercept, F=1964, df=48, P<0.0001) with background wildtype variation.

**Fig. S4.**
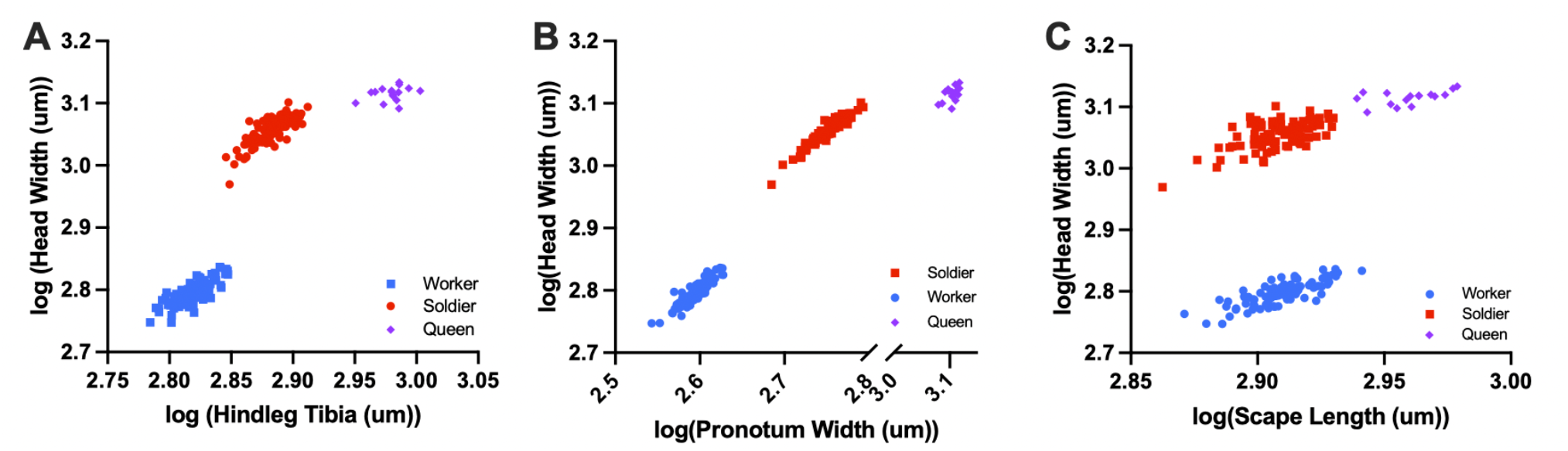
Wild-type variation in *Pheidole dentata* for workers, soldiers, and queens. Log-log plots for **A)** head width to hindleg tibia, **B)** head width to pronotum width, and **C)** head width to scape length.

**Fig. S5.**
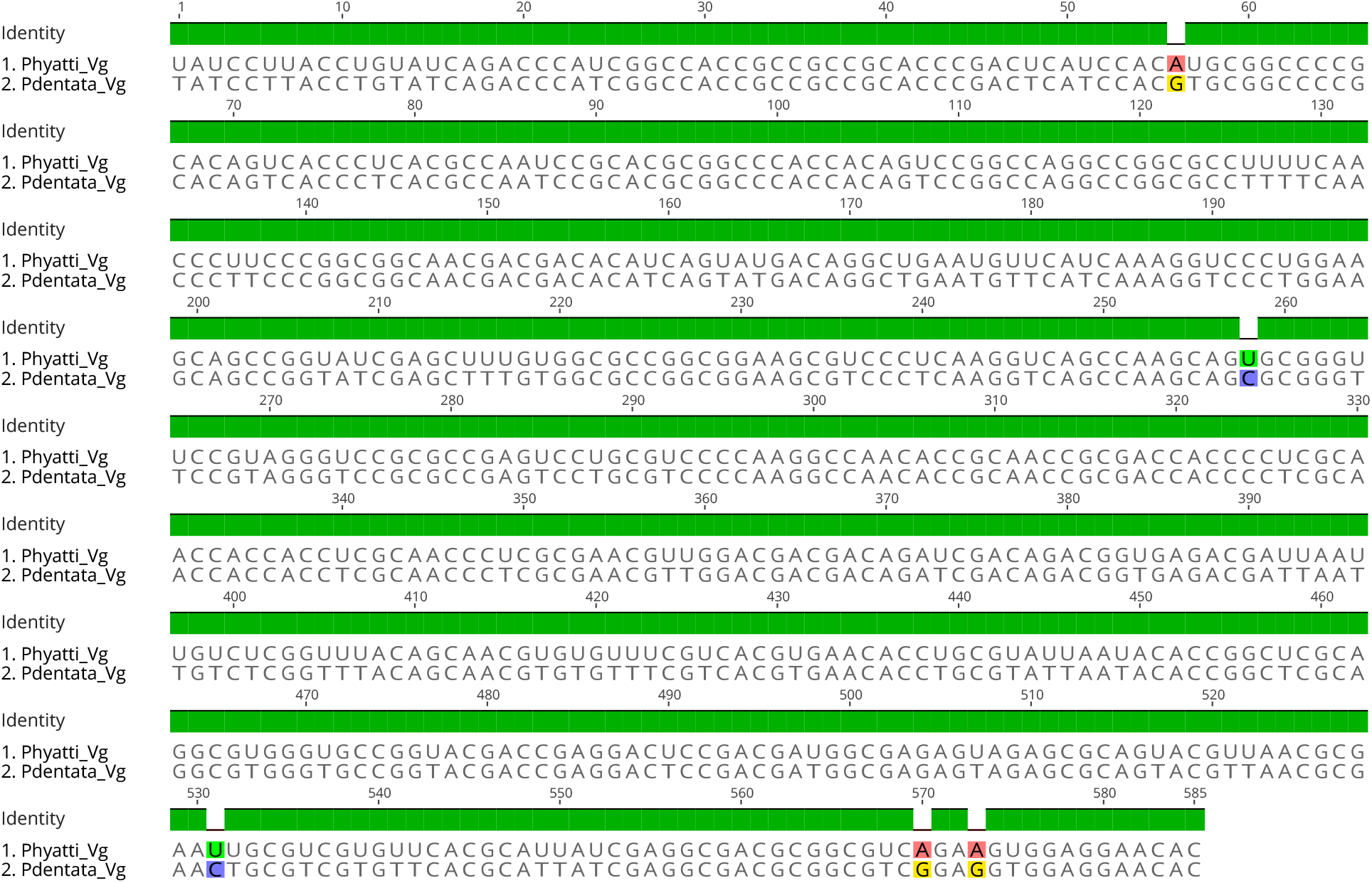
Alignment of the fragment of *vestigial* targeted for RNAi in *P. dentata* as compared to the corresponding *vestigial* target for RNAi in *P. hyatti* from Rajakumar et al., (2018). Sequence is 99.1% identical.

